# Air-interfaced colonization model suggests a commensal-like interaction of *Neisseria meningitidis* with the epithelium, which benefit from colonization by *Streptococcus mitis*

**DOI:** 10.1101/690610

**Authors:** Mathilde Audry, Catherine Robbe-Masselot, Jean-Philippe Barnier, Benoit Gachet, Bruno Saubaméa, Alain Schmitt, Sophia Schönherr-Hellec, Renaud Léonard, Xavier Nassif, Mathieu Coureuil

## Abstract

*Neisseria meningitidis* is an inhabitant of the nasopharynx, from which it is transmitted from person to person or disseminates in the blood and becomes a harmful pathogen. In this work, we addressed the colonization of the nasopharyngeal niche by focusing on the interplay between meningococci and the mucus that lines the mucosa of the host. Using Calu-3 cells grown in air-interfaced culture, we studied the meningococcal colonization of the mucus and the host response. Our results suggested that *N. meningitidis* behaved like commensal bacteria in mucus, without interacting with human cells or actively transmigrating through the cell layer. As such, meningococci did not trigger a strong innate immune response from the Calu-3 cells. Finally, we have shown that this model is suitable for studying interaction of *N. meningitidis* with other bacteria living in the nasopharynx, and that *Streptococcus mitis* but not *Moraxella catarrhalis* can promote meningococcal growth in this model.

## INTRODUCTION

*Neisseria meningitidis* (the meningococcus) is a Gram-negative bacterium that normally resides asymptomatically in the human nasopharynx. For unknown reasons it may cross the epithelial barrier and proliferate in the bloodstream where it becomes one of the most harmful pathogens. *N. meningitidis* effectively adheres to the endothelial cells lining the lumen of blood vessels^1^. From there, bacteria proliferate and cause blood vessel dysfunction^2–6^ responsible for the rapid progression of septic shock, leading in the worst case to *purpura fulminans,* an acute systemic inflammatory response associated with intravascular coagulation and tissue necrosis. *N. meningitidis* can also cross the blood-brain barrier and cause cerebrospinal meningitis^7,8^.

*N. meningitidis* is transmitted from person to person by aerosol droplets produced by breathing, talking, coughing or by direct contact with a contaminated fluid. The natural reservoir of *N. meningitidis* is the human nasopharynx mucosa, located at the back of the nose and above the oropharynx. There, the bacteria encounters a rich microbiota^9–11^ that continuously undergoes changes with age and upon upper respiratory infections^12,13^. The nasopharynx is lined with two main types of epithelium: a pluristratified squamous epithelium that covers 60% of the nasopharynx and a columnar respiratory epithelium^14,15^. In the respiratory tract, cells are protected by a 10-12 μm thick two-layer surface liquid formed by a low-viscosity periciliary liquid (PCL) in contact with the cells and a high-viscosity mucus facing the lumen that retains bacteria, inhaled particles and cell debris (outer mucus)^16,17^ The PCL facilitates ciliary beating that allows effective mucociliary clearance at 6.9 ± 0.7 mm/min^18^. By constantly transporting mucus from the lower respiratory tract to the pharynx from where it is swallowed, this mechanism is considered as the main defense against microorganisms and particles. The mucus layer in which commensal bacteria are restricted is a thick gel formed by mucins and contains many antimicrobial proteins and peptides such as IgA, lysozyme, lactoferrin and human defensins^19–21^. Mucins are a family of at least 22 high molecular weight glycoproteins divided into two classes: membrane associated mucins that are produced by any epithelial cell and gel-forming mucins produced by goblet cells and submucosal glands. In the respiratory tract, the mucus layer is mainly composed of MUC5AC and MUC5B. Their expression is tightly regulated and responds to bacterial infections and to a variety of respiratory tract diseases^17^

The interaction of *N. meningitidis* with epithelial cells has been the subject of numerous studies over the past four decades. However, the means by which meningococci cross the nasopharyngeal wall is still under debate. This may be due to the lack of convenient and relevant model mimicking the nasopharyngeal niche. Most of the previous studies, addressing the adhesion-dependent interaction of *N. meningitidis* with intestinal and respiratory tract epithelial cells, have been performed on cells cultured in liquid media such as RPMI and DMEM. These first studies gave rise to the concept of type IV pili- and/or Opa/Opc-mediated cell colonization. In such a model, meningococci interact with epithelial cells through their type IV pili and form highly proliferative microcolonies that eventually cross the epithelial barrier^22–28^ while Opa and Opc proteins are involved in an active internalization process that is supposed to promote the translocation of bacteria through the cell monolayer^29–31^. Each of these works demonstrated a close interaction of *N. meningitidis* with human epithelial cells. Over the last decade, few studies have focused on the translocation of *N. meningitidis* through the epithelial wall. The work of TC Sutherland in 2010^32^ addresses this question by using Calu-3 human bronchial epithelial cells, a respiratory tract cellular model that can be fully differentiated into a polarized epithelium. Although the authors worked with cells infected in liquid-covered culture (LCC), they proposed to use Calu-3 cells in air-interfaced culture (AIC), a model in which the cells are grown with the apical domain facing the air. They finally concluded that *N. meningitidis* may cross the epithelial layer by the transcellular route using type IV pili. Meanwhile, Barrile *et al* have shown, using Calu-3 cells cultured in LCC, that meningococci may be internalized and transported to the basal domain by subverting the intracellular traffic of the host cells^33^. However, they have also shown that the translocation of bacteria was fully inhibited in highly polarized cells cultured for 18 days.

In addition to these works, a series of *ex vivo* experiments were carried out between 1980 and 1995 using organ cultures instead of immortalized cells^34–37^ The authors have observed a direct interaction between meningococci and explant epithelial cells. This has been associated with the loss of cilia and, for some explants, with the internalization of bacteria in epithelial cells. However, in each of these experiments, the explants were immersed in liquid medium, a protocol that may have altered cell morphology and disrupted the mucus barrier.

The study of meningococcal colonization of the human upper respiratory tract has been hampered by the lack of relevant models. In this work, we took advantage of Calu-3 cells grown under AIC^38^ to study how meningococci colonize the nasopharyngeal niche. Infection of Calu-3 cells revealed the dependence of *N. meningitidis* on mucus in this model. Our results suggest that the mucus protects meningococci against death associated with desiccation and support growth of bacteria. We have shown that the mucus layer sequestered bacteria and that a firm interaction of bacteria with the epithelial cells was rarely observed. Bacteria grew without triggering a strong innate immune response from the Calu-3 cells. Embedded in the mucus, meningococci were protected and fed, expressed less adhesion factors, a high level of iron transporters and type IV pili were not necessary for colonization. Finally, we evaluated the effect of *Streptococcus mitis* and *Moraxella catarrhalis* colonization, two bacteria classically present in the nasopharynx mucosa, on the growth of *N. meningitidis*^39–41^ and showed that co-colonization of *N. meningitidis* with *S. mitis* can promote meningococci growth.

## RESULTS

### Meningococci require mucus production to colonize cells cultured in air-liquid interface

To study the colonization of the human upper respiratory tract by *N. meningitidis,* we used Calu-3 cells cultured on 0.4μm pore membrane under AIC. Cells maintained for a few days in AIC (Week 0) formed a mono-stratified epithelium with only a few spots of mucus on the cell surface (**Figure S1A**). After two weeks of AIC, we observed pseudo-stratified Calu-3 cells covered with a mucus layer. After three weeks under AIC, the epithelium appeared pluristratified with a thick layer of mucus above the cells (**Figure S1A**). Cells cultured for 2 weeks under AIC also revealed the presence of tight junction, microvilli and mucus producing cells (**Figure S1B**). We first considered whether mucus may influence the growth of meningococci on Calu-3 cells cultured under AIC for two days or two weeks (AIC W_0_ and AIC W2, respectively). We added 1.10^6^ meningococci (strain 2C4.3) on the top of cells and assessed epithelial colonization by confocal imaging and quantitative culture (CFU counts), 24 hours after infection. These results were compared to those obtained after infection of Calu-3 cells cultured under liquid-covered culture (LCC). We observed a dramatic decrease in bacterial proliferation 24 hours after infection of AIC W_0_ cultured cells compared to LCC cultured cells (1.35×10^8^ +/− 0.35×10^8^ bacteria per well in LCC; 0.75×10^6^ +/− 0.25×10^6^ bacteria per well in AIC W_0_). This inhibition is less pronounced in AIC W_2_ cultured cells that produced mucus (2×10^7^ +/− 0.64×10^7^ bacteria in AIC W_2_) (**Figure 1A and 1B**). This result suggests that AIC infection is more challenging for meningococci than LCC infection and that the presence of mucus partially compensates for this defect.

**Figure 1:**
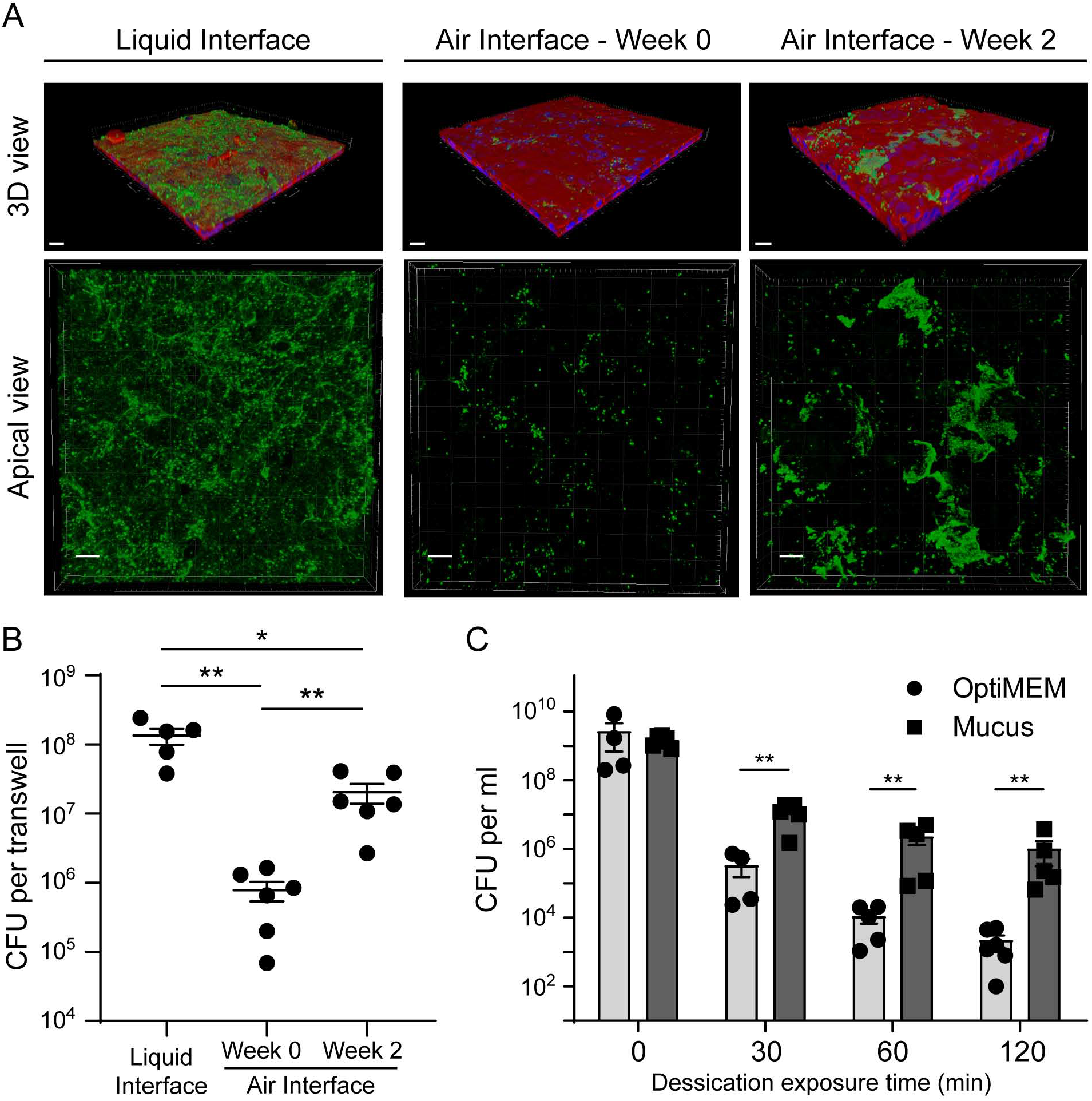
*N. meningitidis* proliferation in an Air Interfaced Culture model. **(A)** Confocal 3D reconstructions showing *N. meningitidis* proliferation, 24 hours after infection of Calu-3 cells. Meningococci were labeled with anti-2C43 antibody (green). Cells were stained with Alexa-conjugated Phalloïdin (red). Nuclei were stained with dapi (blue). Upper panel: 3D view; Lower panel: Apical view. Bar: 20 μm. **(B)** Number of CFU of *N. meningitidis* per well 24 hours after infection (10^6^ bacteria). Data were expressed as mean CFU per filter ± SEM (statistical significance: * *p*<0.05, ** p<0.01; One-way ANOVA; At least five filters; three independent experiments). **(C)** Desiccation assay. Plates were coated with the mucus obtained from Calu-3 cells or with the culture media as control. Bacteria were then grown overnight in these wells containing culture media. The media was then gently removed. Bacteria were dried for 0, 30, 60 or 120 minutes. The number of CFU was then assessed. Data were expressed as mean of CFU per ml ± SEM (statistical significance: ** p<0.01; Student *t* test; At least four wells; two independent experiments).

The mucus layer is known to be a poor nutritive medium that limits the growth of many commensal and pathogenic bacteria. It is a highly hydrated gel that also protects the cell surface from desiccation. We therefore aimed at determining whether the mucus layer could protect *N. meningitidis* against desiccation in an abiotic surface colonization model (**Figure 1C**). We infected plastic wells coated with purified mucus and applied desiccation condition to meningococci for 30, 60 and 120 minutes. Meningococci were particularly sensitive to desiccation. In the uncoated wells (without mucus), the number of living bacteria was reduced by 5.6×10^3^ fold at 30 minutes and 7.9×10^5^ fold at 120 minutes after the beginning of the experiment. Conversely, in mucus-covered wells, the number of living bacteria was reduced by 0.11×10^3^ fold at 30 minutes and 1.46×10^3^ fold at 120 minutes after the beginning of the experiment. Overall, our results indicate that the mucus layer of Calu-3 cells cultured in AIC protect bacteria from desiccation.

### Meningococci are restricted in the mucus layer and do not cross the epithelium

During LCC infection, bacteria readily adhere to human cells and induce host-cell signaling leading to the recruitment of ezrin and actin and the formation of membrane protrusions^42,43^. Conversely, during AIC infection, as it would be the case during colonization of nasopharyngeal mucus, bacteria are deposited on the mucus layer that protects the cells. We therefore studied how *N. meningitidis* interact with the epithelium cultured under AIC. First, we compared the number of CFU recovered from the outer mucus with a fraction containing cells and the cell-attached mucus. The cells were infected with 10^6^ wild type meningococci or its non-piliated derivative (*pilE* mutant; Δ*pilE*), which is unable to adhere to human cells^24^ Up to 80% of wild-type or Δ*pilE* meningococci have been recovered in the outer layer of the mucus, which means that *N. meningitidis* only penetrate slightly through this layer (**Figure 2A**). Interestingly, the same amount of wild-type and Δ*pilE* meningococci was collected in the fraction containing cells and the cell-attached mucus. To better characterize the infection of Calu-3 cells grown using AIC, and determine if the bacteria interacted with the cells, we visualized infected cells by transmission and scanning electron microscopy (TEM and SEM).

**Figure 2:**
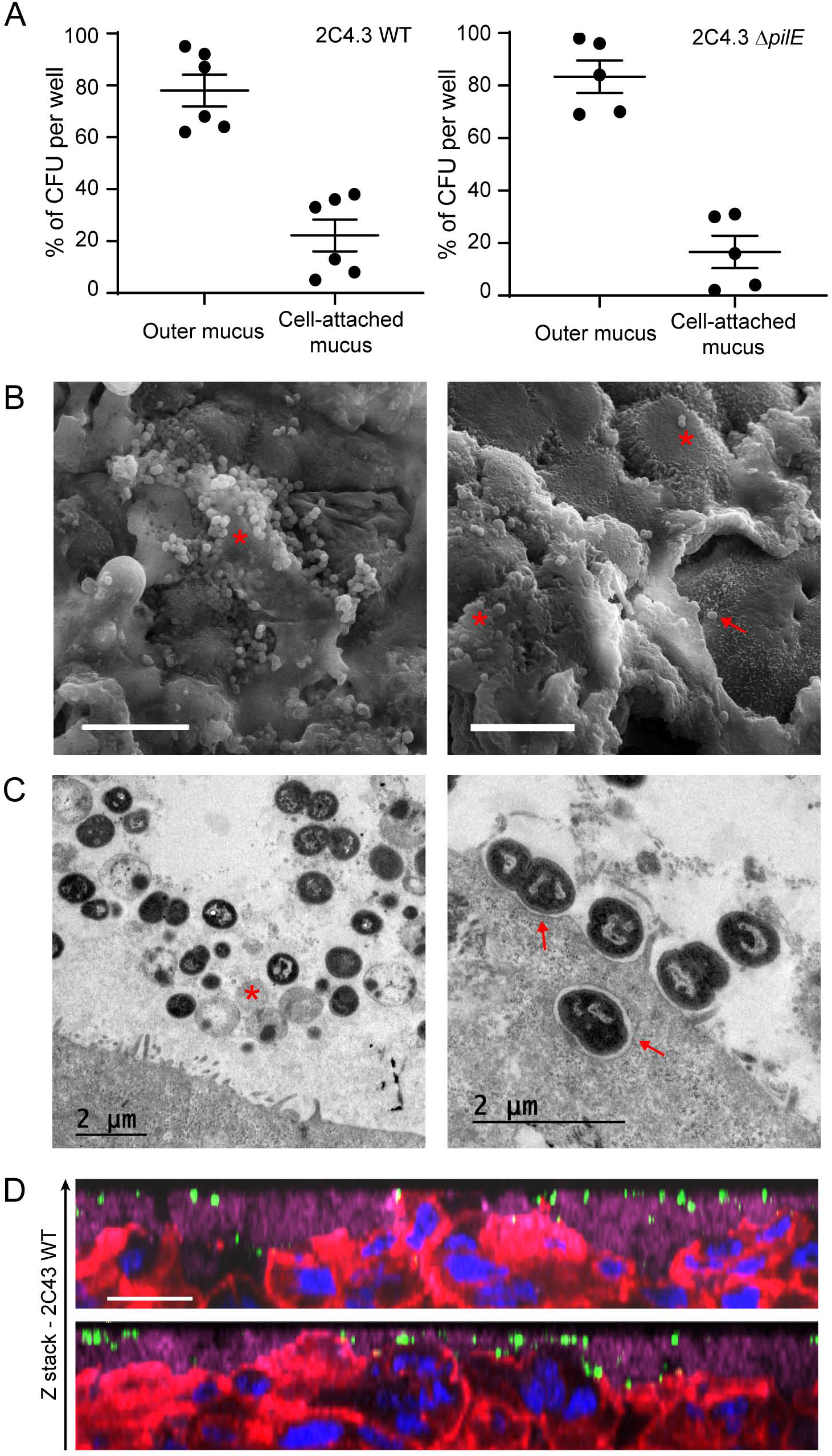
*N. meningitidis* colonize the outer mucus. Calu-3 cells grown in AIC were infected for 24 hours with 10^6^ bacteria. **(A)** After infection, the outer layer of the mucus (outer mucus) was dissociated from the cell-attached mucus using N-acetylcysteine. Bacterial load in the N-acetylcysteine fraction or the cell-attached fraction was determined. Data were expressed as mean percentage of CFU ± SEM (At least five filters; three independent experiments). **(B)** Scanning electron microscopy images showing bacteria trapped in the mucus. *: bacteria; arrow: bacteria that directly interact with Calu-3 cells. Bar: 10 μm. **(C)** Transmission electron microscopy images showing bacteria in the mucus (left) or in contact with cells (right). *: dying bacteria; arrow: bacteria adherent to Calu-3 cells. Bar: 2 μm. **(D)** Z-stack from confocal 3D reconstruction of two different Calu-3 cell layers infected with *N. meningitidis.* Calu-3 cell layers were fixed and immuno-stained with anti-2C4.3 antibody (green) and anti-MUC5AC antibody (purple). Cells were stained with A546-phalloidin (red) and nuclei were stained with dapi (blue). Bar: 20 μm.

We found that most bacteria were trapped in the mucus (**Figure 2B**), and organized into small aggregates of living and dying meningococci, according to cell morphology. Only few bacteria were found in direct contact with Calu-3 cells plasma membrane and we did rarely detect membrane protrusions near bacteria, in contrast to what has been observed previously when cells were infected in LCC^43^. We detected only few internalized bacteria, despite analysis of four different longitudinally cut cell layers. These bacteria were in the vicinity of the apical plasma membrane (**Figure 2D**) or already digested in a vacuole (**Figure S2**). We then studied the infected Calu-3 cells by confocal microscopy. Again, most bacteria were detected in the mucus, stained with anti-MUC5AC antibody. We also observed both by confocal microscopy and TEM, microcolonies of bacteria trapped between epithelial cells in cavities in the upper part of the cell layer (**Figure 3A, B**). However, no bacteria were observed at the basal part of the epithelium as would have been expected if the bacteria had passed through it. In addition, non-piliated Δ*pilE* mutant showed the same spatial location as the wild type strain, suggesting that type IV pili have no role on the location of bacteria in the mucus (**Figure 3A**).

**Figure 3:**
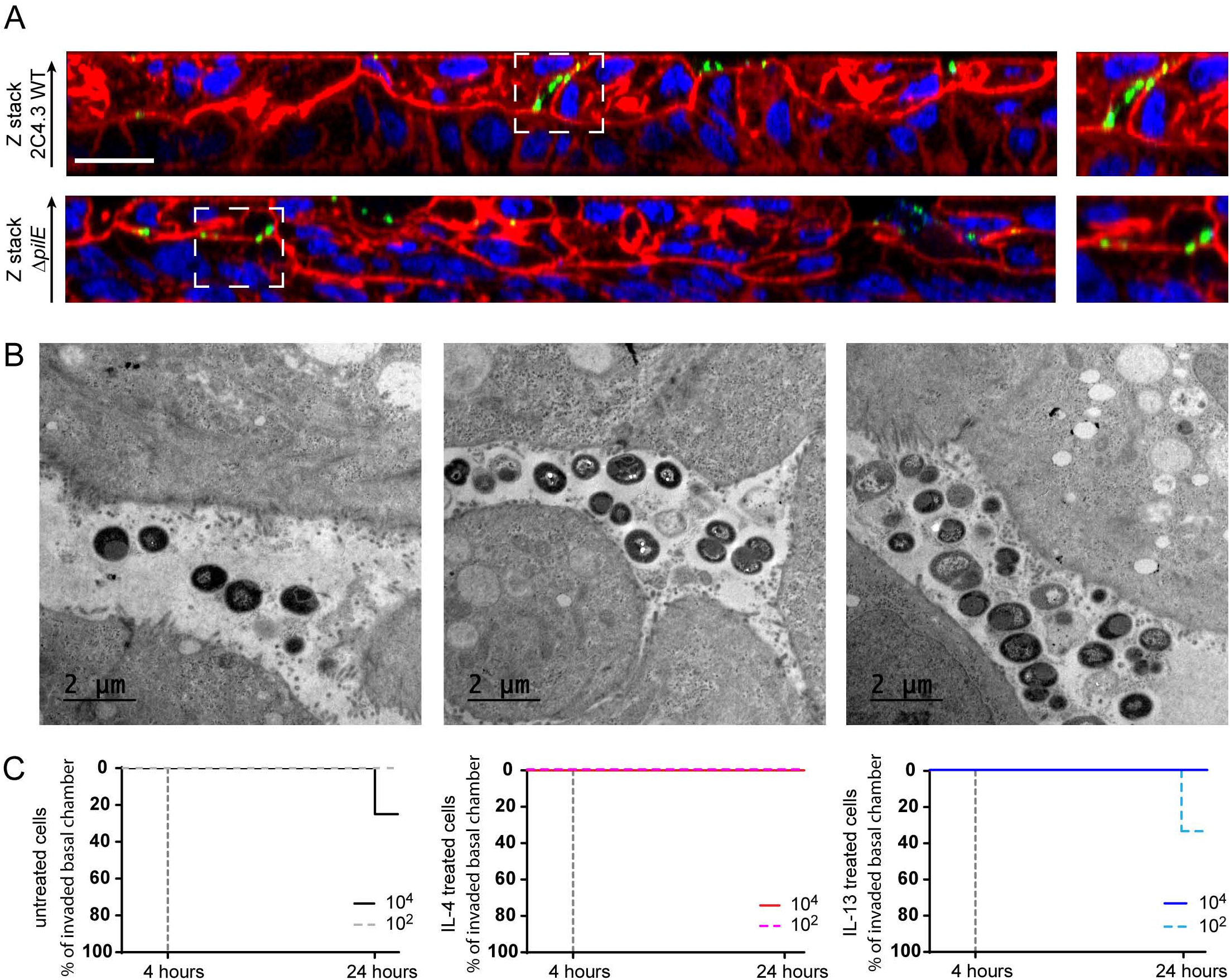
*N. meningitidis* do not cross the epithelial layer. **(A)** Z-stack from confocal 3D reconstruction of Calu-3 cell layer infected 24 hours with 10^6^ bacteria of the wild type strain (2C4.3 WT) or the strain defective for type IV pili (2C4.3 Δ*pilE*). Calu-3 cell layer were fixed and immuno-stained with anti-2C4.3 antibody (green). Cells were stained with Alexa-conjugated phalloidin (red) and nuclei were stained with dapi (blue). Bar: 20 μm. **(B)** Transmission electron microscopy images (longitudinal sections) of Calu-3 cell layer infected for 24 hours with 10^6^ wild type meningococci. Bar: 2 μm. **(C)** Kaplan-Meier plot showing the traversal of wild type *N. meningitidis* across the Calu-3 cell layer. Data were expressed as percentage of invaded basal chamber (untreated cells: 8 filters infected with 10^4^ or 10^2^ bacteria, three independent experiments; IL-4/13 treated cells: 4 filters infected with 10^4^ or 10^2^ bacteria, two independent experiments).

Based on this observation, we studied the translocation of meningococci through the epithelial cell layer. We first grew Calu-3 cells under AIC using 3 μm pore membranes instead of 0.4 μm pore membranes. We choose to infect Calu-3 cells with 10^2^ or 10^4^ bacteria and we first confirmed the proliferation of *N. meningitidis* in these conditions (**Figure S3B, C**). Regardless of the inoculum, the number of colonizing bacteria at 24hrs, 10^7^ CFU, was similar. We then studied the translocation of *N. meningitidis* from the mucus to the basal chamber, by plating the basal media on agar plates, at 4 and 24 hours after infection. We considered a positive translocation when at least one CFU has been recovered from the basal chamber. Interestingly, 24 hours after infection, we detected only 2 out of 16 colonized basal chambers in total, and no bacteria were recovered in the basal chambers 4 hours after infection (**Figure 3C**). Using this model, we then evaluated the effect of IL-4 or IL-13, two cytokines known to be involved in pharyngeal inflammation^44^, on the translocation of *N. meningitidis.* The cells were treated 24 hours with 5 ng/ml IL-4 or IL-13 as previously described^44^ (**Figure 3C**). As expected, the treatment of Calu-3 cells with IL-4 or IL-13 resulted in a 2-fold decrease in TEER (165.9±20.02 Ω.cm2 and 155.3±11.58 Ω.cm2, respectively; **Figure S2A**). However, this has not been associated with an increased traversal of the cell layer by *N. meningitidis* (**Figure 3C**). Our results indicate that meningococci are likely to colonize the outer layer of mucus, from which bacteria can reach the cell-attached mucus but rarely come into contact with cells or cross the epithelial layer cultured in AIC.

### Expression of meningococcal virulence factors during air-liquid infection

Our results suggest that, on mucosa, meningococci are likely to live trapped in the mucus. In these conditions, it is likely that meningococci regulate the expression of their genes differently than in broth. We therefore characterized the relative expression of genes known to be involved in mucosal colonization. We have focused on the expression of genes encoding adhesion factors: *pilE, opaB, opaC* and *nhbA;* genes encoding proteins involved in iron acquisition: *tbpA, lbpA, fetA,* and *tonB; mtrC* encoding the first gene of the *mtrCDE* operon that is involved in drug efflux; *ctrA* that codes for the capsular transport protein A; and the four genes coding for targets of the MenB vaccine, *porA, fhbp, nadA* and *nhbA.* We compared the expression of these genes during AIC infection to their expression during the exponential and stationary phase of growth in broth (**Figure 4**).

**Figure 4:**
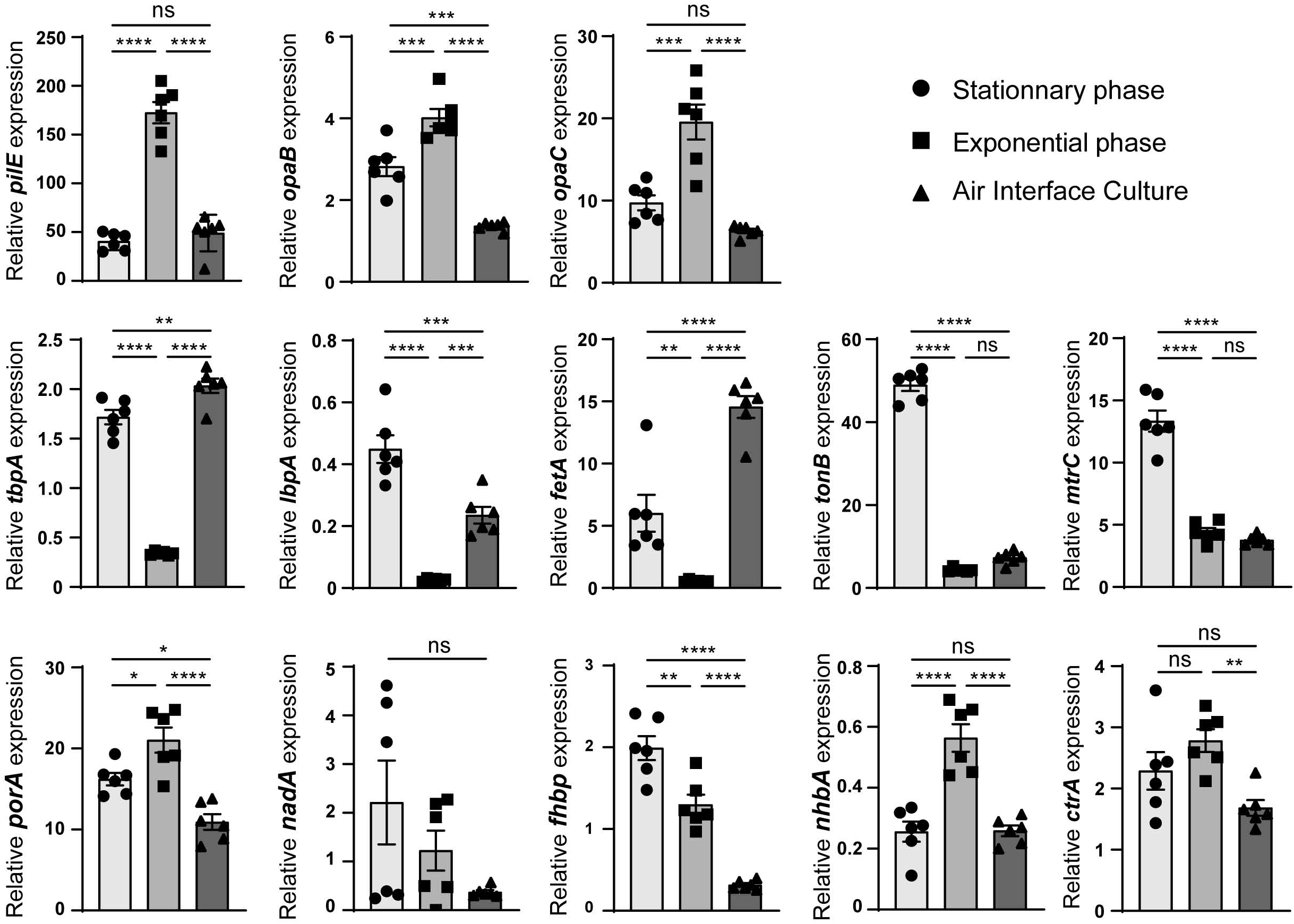
Expression of virulence factors. Total RNA obtained from a 3 hours or 24 hours broth culture or harvested following 24 hours of infection of Calu-3 cells using AIC, was prepared. Gene expression of *pilE, opaB, opaC, tbpA, lbpA, fetA, tonB, mtrC, porA, nadA, fhbp, nhbA and ctrA* was analyzed by quantitative RT-PCR. Gene expression was normalized to that of *pgm* and expressed as relative expression ± SEM (statistical significance: **** p<0.0001; ** p<0.01; * p<0.05; ns: no significant difference; One-way ANOVA; two independent experiment in triplicate).

These genes followed different expression profiles. Expression of the adhesion factors *(pilE, opaB, opaC* and *nhbA)* were comparable between the stationary phase of growth in broth and the infection of Calu-3 cells for 24 hours. In contrast, expression of *pilE, opaB, opaC* and *nhbA* in AIC was decreased with respect to the exponential phase of growth (AIC/exponential phase: 0.28, 0.34, 0.33 and 0.47 fold, respectively). The three iron transporters tested *(tbpA, IbpA, fetA)* were strongly expressed in AIC compared to the exponential phase of growth (AIC/exponential phase: 6.25, 10 and 27 fold, respectively).

It is noteworthy that *fhbp, tonB, mtrC* expression were weak during AIC infection. Their expressions were 6.3, 6.7 and 3.5 fold less expressed in the mucus of Calu-3 cells than in the stationary phase of growth. No major difference in the expression of *nadA,porA* and *ctrA* was observed between the tested conditions. Overall, nine out of the thirteen tested genes appeared to follow the same pattern of expression in AIC than in the stationary phase of growth.

### Meningococci trigger less inflammation in AIC condition

We then addressed the impact of meningococcal colonization on the innate immune response of Calu-3 cells grown in AIC or LCC. We therefore measured the release of ten pro- or antiinflammatory cytokines 24 hours after infection in comparison to the basal release of these cytokines by Calu-3 cells after 24 hours of culture without bacteria (**Figure 5**). After infection we observed that three cytokines were produced in higher amount under LCC infection than AIC infection: IL1β: 2.9 fold increase under AIC versus 15.6 fold increase under LCC; TNFα: 66.4 fold increase under AIC versus 200.4 fold increase under LCC; IL-4: 6 fold increase under AIC versus 31.8 fold increase under LCC. A moderate release of four other cytokines was only detected under LCC: IL-10, IL-13, IL-2 and IL-6 (1.4 fold increase, 1.5 fold increase, 1.6 fold increase and 2.33 fold increase, respectively). Finally, IFNγ secretion was increased by 2 fold under AIC versus 3 fold under LCC, and IL-12 secretion was increased by 3.1 fold under AIC versus 5.1 fold under LCC. Altogether, the pro-inflammatory response assessed by cytokine production, appeared to be higher in the LCC model compared to the AIC model. In addition, during an AIC infection we did not observe any secretion of IL-2, IL-10, IL-13 and IL-6.

**Figure 5:**
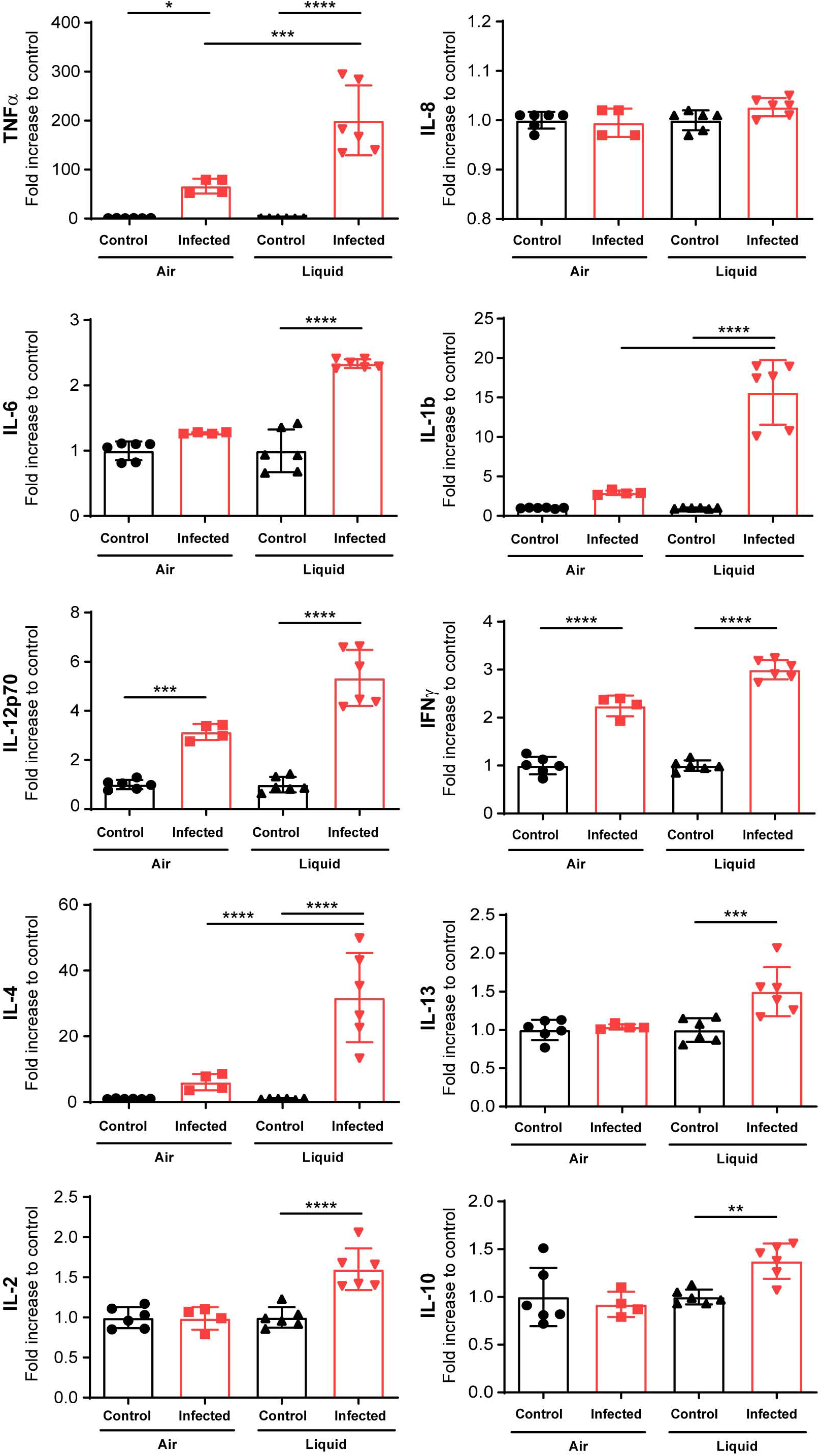
Cytokine expression by infected Calu-3 cells grown in AIC and LCC. Cytokine secretion was investigated in the mucus of non-infected and infected in AIC for 24 hours or in the supernatant of non-infected and infected Calu-3 cells, in LCC for 24 hours. Data were expressed as mean ± SEM of fold increase between infected and non-infected conditions (statistical significance: **** p<0.0001, *** p<0.001, ** p<0.01, * p<0.05; One-way ANOVA: at least two filters read in duplicate).

### *Streptococcus mitis* colonization of Calu-3 cells promotes *Neisseria meningitidis* growth

We next aimed at studying in this model the interplay between *N. meningitidis* and other bacterial species. Among species known to colonize the human nasopharynx, we selected *S. mitis* and *M. catarrhalis* as representative bacteria^39–41^. We first infected Calu-3 cells, cultured in AIC on a 0.4μm pore membrane, with 1.10^5^ *S. mitis* or *M. catarrhalis.* Both were able to survive in the mucus of Calu-3 cells, but we did not detect proliferation 48 hours after infection (*S. mitis*, inoculum: 3.08×10^5^±0.8 10^5^, 48 hours: 2.7×10^5^±1.01; *M. catarrhalis,* inoculum: 4.12×10^5^±1.9 10^5^, 48 hours: 6.07×10^5^±2.38). We then infected Calu-3 cells colonized by *S. mitis* or *M. catarrhalis* with 1×10^6^ meningococci (wild-type 2C4.3 strain) for 24 hours (**Figure 6**). As control, naïve uninfected Calu-3 cells were infected by *N. meningitidis*. Our results showed that the co-infection with *S. mitis* significantly improved meningococcal colonization while *M. catarrhalis* had no effect (**Figure 6**). Interestingly, the positive effect of the *S. mitis/N. meningitidis* co-infection on the growth of meningococci appeared to be specific of the AIC model. A 24 hours broth co-culture of 1.10^5^ *S. mitis* and 1.10^6^ *N. meningitidis* revealed a slight decreased in meningococcal growth (**Figure S4A**). Overall, these results support the hypothesis that *N. meningitidis* growth in AIC may be facilitated by other bacteria of the nasopharyngeal niche.

**Figure 6:**
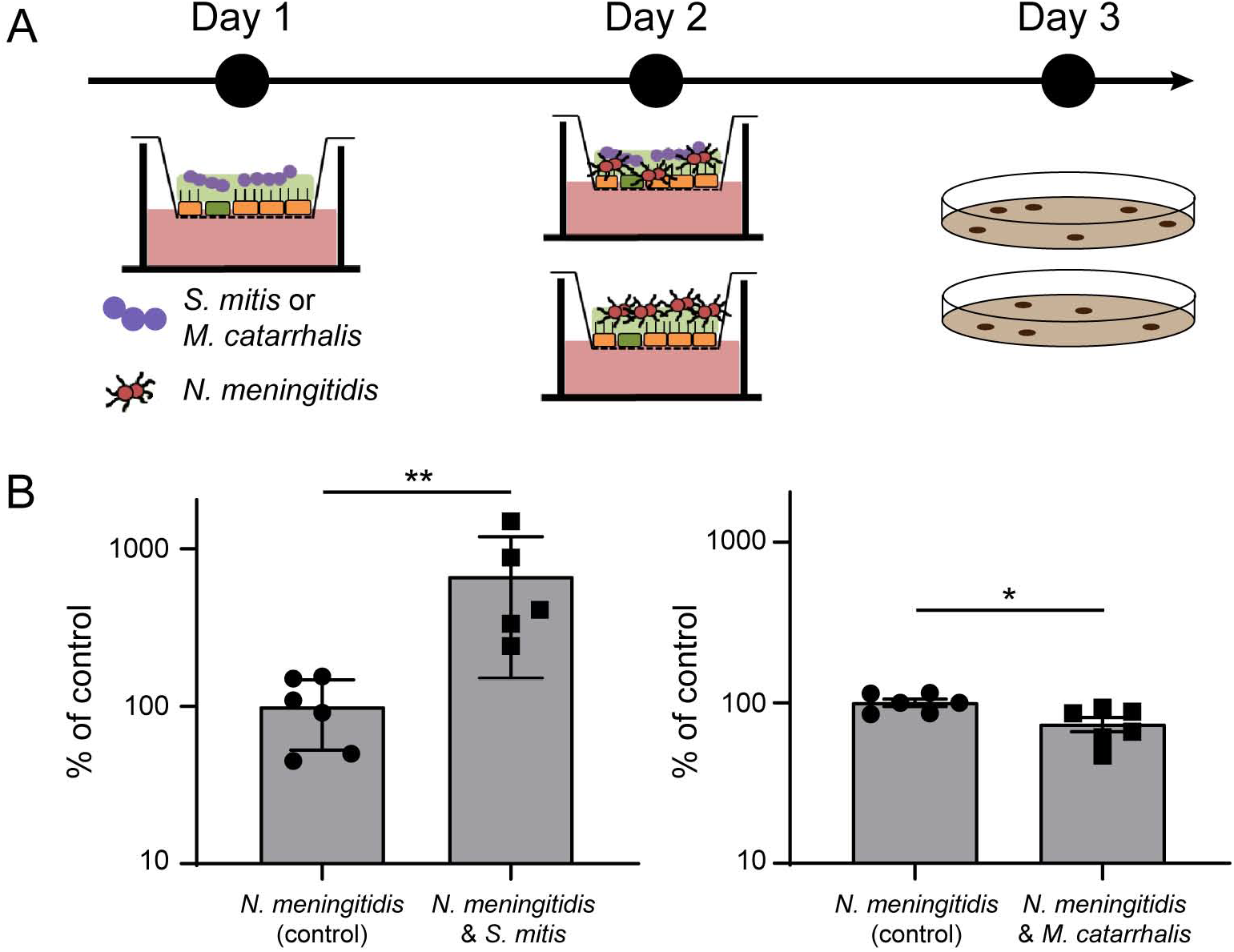
Co-infection with *S. mitis* and *M. catarrhalis.* **(A)** Schematic representation of the protocol followed for co-infection. **(B)** At day 1, cells were infected with 10^5^ bacteria (*S. mitis* or *M. catarrhalis*). At day 2, cells were then infected with 10^6^ meningococci. At day 3, bacteria were collected and CFU were determined. CFU of meningococci after 24 hours of co-culture were expressed as mean percentage of the control experiment ± SEM (CFU of meningococci in mono-culture) (statistical significance: ** p<0.01, * p<0.05; Student *t* test; At least five filters, three independent experiments).

Unlike *M. catarrhalis, S. mitis* is able to hydrolyze glycans, which are very abundant on mucin proteins^45^. The hydrolysis of mucins’ carbohydrates might thus provide an additional source of carbon and nutrient for meningococci, which could be responsible for a significant increase in meningococcal colonization. We investigated mucins’ glycosylation profiles by mass spectrometry after infection or co-infection of Calu-3 cells with *S. mitis* and *N. meningitidis* (**Table 1 and S1**). We observed a moderate sialylation of mucins after meningococcal colonization, since 59% of the oligosaccharides detected were sialylated compared to 36.2% in non-infected cells. The glycosylation profile of mucins has changed dramatically after infection with *S. mitis* (**Table 1 and S1**). We noticed a complete release of sialic acid from the mucins and a global simplification of the O-glycans, while infection with inactivated streptococci did not alter the glycosylation profile of mucins (**Table S1**). This confirms that *S. mitis* is capable of hydrolyzing mucins’ O-glycans. A process that correlates with an increased growth of *N. meningitidis*.

**Table 1.**
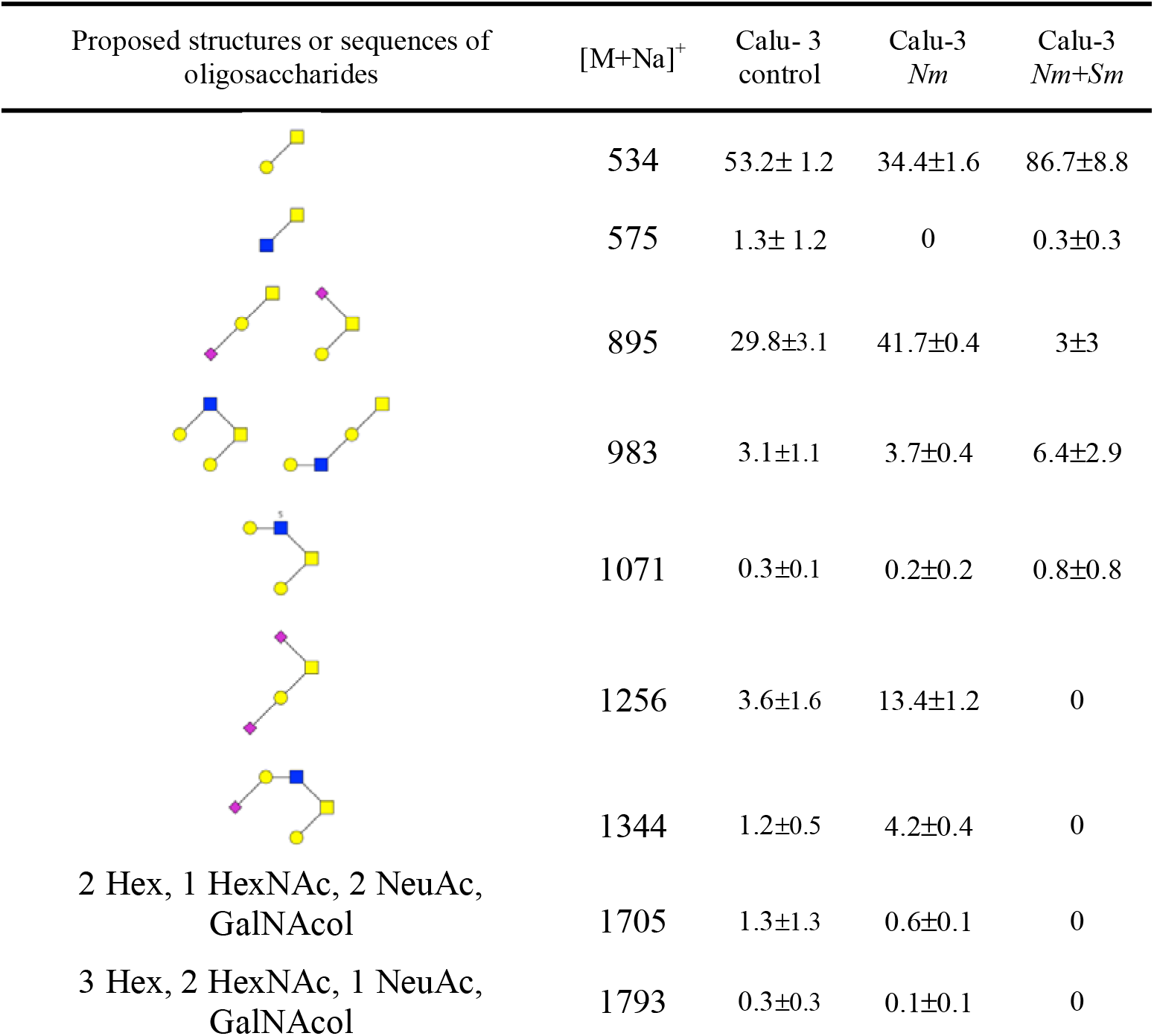
Highlights on sialylated oligosaccharides, and their non sialylated form, identified on Calu-3 mucins, before (control) and after infection with *N. meningitidis (Nm)* or co-infection with *N. meningitidis* and *S. mitis* (*Nm+Sm*). The relative percentage of each oligosaccharide was calculated based on the integration of peaks on MS spectra. Two independent experiments of 5 different filters were studied in bulk. Results are presented as the mean relative percentage of each oligosaccharide ± SEM.

| Proposed structures or sequences of oligosaccharides | [M+Na]<sup>+</sup> | Calu- 3 control | Calu-3 *Nm* | Calu-3 *Nm+Sm* |
| --- | --- | --- | --- | --- |
| | 534 | 53.2± 1.2 | 34.4±1.6 | 86.7±8.8 |
| | 575 | 1.3± 1.2 | 0 | 0.3±0.3 |
| | 895 | 29.8±3.1 | 41.7±0.4 | 3±3 |
| | 983 | 3.1±1.1 | 3.7±0.4 | 6.4±2.9 |
| | 1071 | 0.3±0.1 | 0.2±0.2 | 0.8±0.8 |
| | 1256 | 3.6±1.6 | 13.4±1.2 | 0 |
| | 1344 | 1.2±0.5 | 4.2±0.4 | 0 |
| 2 Hex, 1 HexNAc, 2 NeuAc, GalNAcol | 1705 | 1.3±1.3 | 0.6±0.1 | 0 |
| 3 Hex, 2 HexNAc, 1 NeuAc, GalNAcol | 1793 | 0.3±0.3 | 0.1±0.1 | 0 |

## DISCUSSION

In this work, we adapted an experimental model, based on Calu-3 cells cultured in AIC, to study the behavior of *N. meningitidis* in the mucus of the respiratory tract. We have shown that meningococci are trapped in the mucus layer where bacteria are protected from desiccation and are likely to find nutrients to grow. We found no evidence of an active passage of *N. meningitidis* through the epithelial layer and we observed that type IV pili were not important for growth or motility/mobility in this model. Similarly, we showed that virulence factors were poorly expressed in this model compared to culture in broth. Strikingly, this suggests commensal-like behavior of *N. meningitidis,* a hypothesis that is supported by the poor cytokine response observed 24 hours after infection. We took advantage of this model to investigate the effect of other bacteria on the growth of *N. meningitidis*. We have shown that *S. mitis,* which is able to hydrolyze glycans, facilitates the growth of meningococci in a co-infection protocol.

Most studies aimed at determining the behavior of meningococci on epithelial cells has been done with cells that have been cultured and infected in LCC. Although these investigations provided a comprehensive description of the interaction between *N. meningitidis* and human epithelial cells, the scientific community has not been able to conclude on the question of how and when meningococci cross the nasopharyngeal epithelium. During LCC infections, bacteria easily proliferate in the cell culture medium that contains amino acids, carbon source and protein extracts. This allows *N. meningitidis* to grow and eventually cover almost completely the Calu-3 cell layer. When we infected Calu-3 cells cultured in AIC for 2 weeks, we observed a 6 fold decrease in the total number of CFU per filter 24 hours after infection. The bacteria were mainly found in the mucus where they organized into small groups of living and dying meningococci. As a consequence, bacteria rarely interact with human cells and we have barely found *N. meningitidis* in direct contact with the plasma membrane of these cells. In view of this result, we asked if *N. meningitidis* can cross the epithelial layer grew in AIC. After infection of cells cultured on 3 μm pore membranes, we detected bacteria in the basal chamber of only 2 out of 8 wells for the highest inoculum (10^4^ CFU) and 0 contaminated chambers for the lowest (10^2^ CFU). We then treated Calu-3 cells with IL-4 or IL-13 for 24 hours, two cytokines that induce TEER decrease and induce mucus production^44^ As expected, these two cytokines led to a reduction in TEER. But, this has not been followed by an increase in the translocation of bacteria through the epithelial layer. All these results suggested that the traversal observed in this experiment was only stochastic and probably due to the heterogeneity of the mucus layer on the surface of the wells. To support this hypothesis, we have never observed any bacteria, inside or outside the cells, and in the vicinity of the porous membrane.

These results suggest that *N. meningitidis* growing in the mucus of epithelial cells did not alter the epithelial layer during the course of the experiment. We therefore studied the production of cytokines by epithelial cells grown in AIC or in LLC. Strikingly, we observed that three major inflammatory cytokines (IL-6, TNFα and IL-1β) were produced less during infection in AIC than in LCC. The two anti-inflammatory cytokine IL-10 and IL-4 were also produced less after infection, suggesting an overall reduction in the cytokine response during infection in AIC. However, after infection, Calu-3 cells secreted IL-12 and IFNγ to the same extend whether infection was in AIC or LCC. IFNγ and IL-12 are known to be associated with macrophage and dendritic cells response, although there are evidences that epithelial cells produce these cytokines after infection with microbes^46,47^ IFNγ has pleiotropic effects on the epithelial cells of the respiratory tract. This cytokine has been shown to reduce MUC5AC expression, which may lead to a decrease in the barrier property of the respiratory mucus^48^.

In the meantime IFNγ induces the expression of CEACAM receptors^49^ that are the receptors for the meningococcal adhesin Opa, known to be involved in the internalization of bacteria. Conversely, IFNγ may promote the barrier function of lung epithelial cells^44^ Finally, we did not detect IL-8 secretion after infection of Calu-3 cells. In our cytokine assay, AIC cultured cells were generally less reactive than cells that have been cultured in LCC, indicating that the mucus layer is likely to protect cells and retain pathogen-associated molecular patterns, resulting in a reduction in the innate immune response. However, our model, which did not include immune cells, only gave a partial overview of the innate immune response.

We observed that the expression of virulence factors by *N. meningitidis* varied accordingly to growth status and that three virulence genes *pilE, mtrC* and *fhbp* were significantly silenced during infection in AIC. The expression of *pilE* during infection of the AIC model appeared to be similar to that observed during the stationary phase of growth in liquid culture. This was correlated with the absence of a role for type IV pili during the colonization of Calu-3 cells. Although most of the meningococcal strains found *in vivo* were piliated, the role of type IV pili during growth in the mucus was probably not related to motility and/or interaction with epithelial cells. However, we could not exclude an interaction between type IV pili and mucins. Conversely, the expression of *fhbp* during infection in AIC was dramatically decreased compared to the exponential and stationary phase of growth in broth. This suggests that mucus is sensed by bacteria and in which *N. meningitidis* will adapt the expression of its virulence genes. Interestingly, fHBP is a key virulence factor of *N. meningitidis*^50^ necessary for binding to human factor H and that inhibits the host alternative complement pathway. The role of fHBP in the respiratory tracts is not clear. While the respiratory mucus contains complement components^51,52^, the bactericidal activity itself of the complement is not clearly defined against *N. meningitidis*. For instance, acapsulated strains are regularly recovered by swabbing whereas an active component system should have eliminated these strains. It was therefore not surprising to observe the lack of regulation of *ctrA* gene expression between the different conditions tested. Based on our results, and in the context of the MenB vaccine, it may be important to further investigate the expression of *fhbp* in the context of respiratory mucus. Finally we showed that *lbpA, tbpA* and *FetA* were highly expressed in AIC, confirming the low concentration of free iron in this model.

The nasopharynx is colonized by six main genera: *Haemophilus, Streptococcus, Moraxella, Staphylococcus, Alloiococcus and Corynebacterium*^40,41^. The impact of the microbiota on the growth, survival and expression of *N. meningitidis* virulence factor is not yet known. Here, we have used the AIC model to address the impact, on *N. meningitidis* growth, of the colonization by two of these bacteria. We infected Calu-3 cells with *S. mitis* or *M. catarrhalis.* Interestingly, we observed that *S. mitis* promoted meningococcal growth 24 hours after infection. This result was not expected since it is known that the pyruvate oxidase (SpxB) of *Streptococcaceae* produces a high amount of hydrogen peroxide and inhibits the growth of *N. meningitidis* in broth^53^. Okahashi has shown that *S. mitis* also expresses SpxB, which may be deleterious for Calu-3 cells^54^. We therefore studied the growth of co-cultured meningococci with *S. mitis* in broth (Figure S4). As described, a high ratio of *S. mitis* killed meningococci, while a ratio of 1 *S. mitis* per 10 *N. meningitidis* is sufficient to reduce the total number of meningococci by two after one day of co-culture. Conversely, in AIC, *S. mitis* promotes the growth of *N. meningitidis,* suggesting that *S. mitis* was less active against meningococci in AIC conditions. In addition, our glycomic analysis indicated that *S. mitis* is capable of hydrolyzing mucin O-glycans while *N. meningitidis* cannot. This was expected since *S. mitis* is known to express many glycosyl hydrolases^45^. Since sialic acids were released from the O-glycans, we assessed whether this could provide a growth advantage for *N. meningitidis.* As anticipated, meningococci were unable to grow in the presence of sialic acid as sole carbon source in broth, and the addition of sialic acid in the mucus of Calu-3 cells was not sufficient to enhance the growth of meningococci (data not shown). However, it can be hypothesized that *S. mitis* might increase the concentration of other nutrients that may be metabolized by *N. meningitidis*.

All together, our results have shown that infection of mucus-producing cells in AIC is different from that of conventional experiments performed in LCC. While these latter works have investigated the interaction of *N. meningitidis* with epithelial cells, which is likely to occur after substantial inflammation or mechanical breach in the mucus layer, our present study emphasizes that *N. meningitidis* is certainly trapped in the mucus layer and rarely interacts with human cells while the host response is less pronounced. Further work will be needed to better understand how *N. meningitidis* regulates its virulence factors and cohabit with other bacterial species in the mucus.

## MATERIALS AND METHODS

### Bacterial strains and growth conditions

*N. meningitidis* NEM 8013 (2C4.3), a piliated capsulated serogroup C strain, and its isogenic non-adhesive PilE defective mutant (Δ*pilE*) were used in this study^27^

*Streptococcus mitis* B26E10 (referenced as 0902 230473 in Necker Hospital collection) and *Moraxella catarrhalis* B18F4 strains (respectively referenced as B18F4 in Necker Hospital collection) were isolated from a patient in the Necker hospital (Paris). Except for *Streptococcus mitis* strain grown on chocolate agar polyvitex plates, all strains were grown on BHI-agar plates supplemented with 5% horse serum at 37°C in a 5% CO_2_ incubator. Antibiotics were used at the following concentrations: kanamycin at 100 μg/ml, vancomycin at 20 μg/ml, polymyxin at 15 μg/ml.

### Cell culture

Calu-3 epithelial cells (ATCC HTB-55) were maintained in optiMEM medium (Life Technologies) supplemented with 5% fetal bovine serum, HEPES, minimum amino acids solution and penicillin-streptomycin antibiotics. Cells were grown in a 5% CO_2_ incubator at 37°C. Cells were grown on polyester 0.4 μm pore membrane cell culture filter (Corning, Transwell®). For traversal assays, 3 μm pore membrane were used. Prior to cell seeding, filter’s membranes were coated with type IV human placenta collagen (Sigma) for 24 hours. Cells (3.10^5^) were seeded onto the apical side of membranes and were maintained in 200 μl of culture medium in the apical chamber and 1.2 ml in the basal chamber. In AIC conditions, the apical culture media was removed after five days and cells were allowed to grow at air-interface for 3 to 6 days (week 0) or for 14 to 17 days (week 2). Liquid-covered layers were seeded and cultured as described above, except that the apical media were maintained all along. The transepithelial electrical resistance across air interfaced culture was measured with a Millicell® Voltohmmeter (Millipore). Notably, the barrier function of the Calu-3 cell layer that have been grown on a 3 μm pore membrane were decreased, as indicated by measurement of the transepithelial electrical resistance (TEER: 357±19.83 Ω/cm^2^ using 0.4 μm pore membrane; 258±14.63 Ω/cm^2^ using 3 μm pore membrane) (**Figure S3A**).

### Infection

#### Infection with N. meningitidis

Two days before infection, antibiotics were removed from the culture media. On the day of infection, a suspension of bacteria from an overnight culture on agar plate was diluted to a bacterial concentration of 5.10^7^ CFU/ml and cultured for 2 hours at 37°C in optiMEM medium. The air-interfaced culture cells were infected on the apical side with a 10 μl of a bacterial suspension containing 10^6^ CFU per 10μl unless specified. The next day, cells were collected by scratching and thoroughly vortexed, then CFU were counted by plating serial dilutions onto agar plates. The same protocol was applied for liquid interface infections, except that a volume of 200 μl of bacterial suspension was used (10^6^ CFU per 200 μl). In order to separate bacteria present in the outer mucus fraction or in cell-attached mucus fraction, an optiMEM-0.1% N-acetylcysteine solution was incubated for 15 minutes on top of the cells and harvested. This process was repeated 3 times and CFU were counted in this outer mucus fraction by plating serial dilutions. Then for cell-attached mucus fraction, scratching collected cells were vortexed and bacterial loads were assessed by plating CFU.

#### Transmigration assay

One day prior infection with bacteria, human interleukine-4 (IL-4) and interleukine-13 (IL-13) were added in the basal chamber of Calu-3 cells cultured in AIC at 5 ng/ml each. Media were replaced immediately before infection and IL-4 or IL-13 were added freshly. At 4 and 24 hours post-infection, media from the basal chamber of untreated or treated cells were collected and centrifugated. Pellets were resuspended in 200 μl and serial dilutions were cultured on agar plates.

#### Co-infection and co-culture

At day-0, either *Streptococcus mitis* or *Moraxella catarrhalis* strains grown on agar plates overnight were resuspended in optiMEM medium and cultured in optiMEM medium for 2 to 3 hours at 37°C. After reaching the exponential growth phase, Calu-3 cells grown for two weeks in AIC were infected with 10μl of bacterial suspension containing 1.10^5^ CFU. For the control filters, 10 μl of sterile medium was added on top of cells. At day-1, control and infected filters were infected with 1.10^6^ of *N. meningitidis* 2C4.3 strain as previously described. At day-2, bacteria were harvested by scratching and cultured on selective medium agar plates. The 2C4.3 strain was selected on vancomycin (20 μg/ml) when co-cultured with *S. mitis* and on polymyxin (15 μg/ml) when co-cultured with *M. catarrhalis.* During the assay, the cells were incubated at 37°C in a 5% CO_2_ incubator.

### Immunofluorescence assay

#### Fixed cells

For immunofluorescence assays, Calu-3 cells were grown in AIC and infected for 24 hours. The filters were fixed with 4% paraformaldehyde for 1 hour at room temperature, washed two times with PBS and permeabilized for 10 minutes with PBS-0.1% X-100 Triton and 10 minutes in PBS-0.1%BSA 0,1% X-100 Triton (staining buffer). The cells were then incubated with an anti-*N. meningitidis* strain 2C4.3 (anti 2C4.3) (and an anti-MUC5AC monoclonal antibody (clone 45M1; life technologies) in staining buffer for 2 hours. After three washes in PBS, the filters were incubated with Alexa-conjugated secondary antibodies for 2 hours. Nuclei DNA and actin were respectively stained with DAPI at 1 μg/mL and Alexa-conjugated phalloidin (Invitrogen). After several washes, the membranes were cut from the plastic support and the coverslips were mounted in Mowiol for observation.

#### Living cells

Because the mucus could not be easily preserved through fixation, its production over time was monitored by imaging living cells labeled with an Alexa-conjugated Dextran at 1 mg/ml (MW 10000, Life technologies) and Cell Trace Calcein Red Orange AM at 2.5 μM (Life technologies) was used to stain the epithelium. The cell tracer was added in the basal chamber for one hour while the dextran solution was added on top of cells. Both solutions were removed and washed before confocal acquisition. During acquisition, the cells were maintained at 37°C under 5% CO_2_.

### Image analysis

For three-dimensional reconstruction, image acquisition was performed on a laser-scanning confocal microscope (Leica TCS SP5). Fluorescence microscopy images were collected and processed using the Leica Application Suite AF Lite software. Each channel was adjusted for better visualization. 3D reconstruction, z-stack pictures and post-treatment analyses were performed using Imaris software.

### Electron microscopy

#### Chemicals

Crystalline osmium tetroxide (OsO4), sodium cacodylate, 25% glutaraldehyde and Epon were from Euromedex (Souffelweyersheim, France). Hexamethyldisilazane (HMDS) was from Sigma-Aldrich (Lyon, France). Perfluoro-compound FC-72 was from Fisher Scientific (Illkirch, France).

#### Electron microscopy

All incubations were performed at room temperature. Whole inserts were fixed in 1% OsO4 diluted in FC-72 for 90 minutes, rinsed in FC-72 for 30 min and fixed in 2.5% glutaraldehyde diluted in 0.1 M sodium cacodylate buffer, pH 7.4 for 90 minutes. The inserts were then rinsed in cacodylate buffer (2×30 minutes) and immersed in an ascending concentration of ethanol solutions (30%, 50%, 70%, 95%, 100%, 100%, 100% – 10 minutes each) for dehydration. For SEM, dehydration was completed with HMDS/ethanol (1/1, v/v) for 10 minutes and HMDS for 10 minutes. After overnight air drying, each filter was removed from the insert using a small scalpel blade, placed on a double-sided sticky tape on the top of an aluminum stub and sputter coated with Au/Pd. Images were acquired using a Jeol LV6510 (Jeol, Croissy-sur-Seine, France). For TEM, inclusion was performed by immersing the inserts in a Epon/ethanol mixture of increasing Epon concentration (1/3 for 60 minutes, 1/1 for 60 minutes, 3/1 overnight) and finally in pure Epon (two changes in 48 hours). Each filter was removed from the insert and placed in an embedding capsule, with cells facing down. After resin polymerisation (2h at 37°C then 72h at 60°C), the block was sectioned so as to produce section of the cell layer. Ultrathin sections (80 nm) were stained in lead citrate and examined in a Jeol 100S (Jeol, Croissy-sur-Seine, France) at an accelerating voltage of 80 kV. Living bacteria were defined as circular cells and electron dense.

### Desiccation assay

Plastic wells were coated with optiMEM as a control or with mucus overnight. The mucus was extracted from Calu-3 cultured in AIC for two weeks using 0.2% β-mercaptoethanol in optiMEM (3 washes of 20 minutes each). For infection, 5.10^6^ bacteria in 100 μl of media were added on top of the coated plastic well and incubated at 37°C under 5% CO_2_ overnight. The next day, culture supernatants were gently removed and sedimented bacteria were exposed to desiccation for 0, 30, 60 and 120 minutes. Bacteria were harvested and enumerated by quantitative culture into agar plates.

### Quantitative RT-PCR

#### RNA isolation

Total RNA was isolated from *N. meningitidis* cultured at 37°C in optiMEM medium for 3 hours (exponential growth phase), overnight (stationary phase), or from infected cells (AIC) after 24 hours. In those three conditions, bacteria were pelleted by centrifugation at max speed in a microcentrifuge for 2 minutes and quickly resuspended in cold Trizol solution.

Samples were frozen and stored at −80°C. Samples were then treated with chloroform and the aqueous phase was collected and used in the RNeasy Clean-up protocol (Qiagen). RNA samples were incubated with turbo DNase (Invitrogen) for 1 hour at 37°C before cleaning-up on RNeasy mini-column. Elution of RNA was done in nuclease-free water and 1 μl of rRNasin (Promega) was added before storage.

#### Retrotranscription

cDNA synthesis reactions were carried out using the Lunascript RT Supermix kit (NEB) and 500 ng of RNA was used for each reaction.

Quantitative RT-PCR: The 20 μl reaction consisted of 10 μl Luna Universal qPCR Master Mix, 0.5 μl 10 μM of each primer, 1 μl of cDNA and 8 μl of nuclease-free water. Pairs of primers were designed with Primer3Plus software *(pilE* forward TTTGCGACTGTAACGCTTTG, reverse GCCATCCTTTTGGCTGAAGG; *porA* forward TCCCTTGAAAAACCATCAGG, reverse CAATTTCGGTCGTACTGTTT; *<u>nadA</u>* forward AAATTAGAAGCCGTGGCTGA, reverse TGCAGCGACAGCTTCGGCCT; *fhbp* forward CATACCGCCTTCAACCAACT, reverse GTTCGGCGGCGGCAAGCTCG, *nhbA* forward AAACGCCATTAGCCACATTC, reverse CCACGGCACCGAATATGCCA; *pgm* forward GCGAAGCCATAATGGAAAAA, reverse CTTTGCGGCAGGTTGTTTAA) *TonB* forward TCAGCAGCCTAAGGAAGAGC, reverse CTGCCTTCTCCGCGCCCCGT; *LbpA* forward GATAAGGCGGTGTTGTCGTT, reverse TGGAAGCATCGTAACCGAAG; *TbpA* forward GCAGTGGGGGATTCAGAGTA, reverse GGATGGGTATCCTCAACCGG; *FetA* forward CGGCAGAAAATAATGCCAAT, reverse GTGCCGTTGCCGCCGCCGAA; *ctrA* forward GTTTGGCGATGGTTATGCTT, reverse CGGCCTTTAATAATTTCCTG; *mtrC* forward CCGTACCGAACGATCAGAAT, reverse CTTCAGACCCGACGTAACAA; *OpaB* forward CTTGTCCGCCATTTACGATT, reverse TGATACAAGCTTGCCTGGCT; *OpaC* forward AACATCCGTACGCATTCCAT, reverse AAGCCGAGCGAGGAGACGGC. Gene expression levels were normalized by that of the housekeeping gene *pgm* (NMV_1606). Appropriate no-RT controls were carried out to ensure accuracy of the results.

### Cytokine quantification assay

#### Sample preparations

Calu-3 cells were grown on transwell either in AIC or LCC during two weeks. For AIC, cells were incubated at 37°C for 20 minutes with 100 μl of Ringer solution and that apical supernatants were collected. This step was repeated once, and samples were kept on ice. For liquid interfaced culture, the 100 μl of medium in the apical chamber was collected and cells were washed with another volume of 100μl of optiMEM medium. All samples were vortexed and centrifugated at 4°C for 5 minutes at maximum speed to eliminate bacteria and debris. Supernatants were harvested, snap-freeze in liquid nitrogen and kept at −80°C before processing.

#### Cytokines measurement

cytokines in cell supernatants were quantified by electrochemiluminescence multiplex assay kits from Meso Scale Discovery (Rockville, MD, USA). Briefly, 25 μl of supernatant were added to 96-well multi-spot plates and the assays were performed following the manufacturer’s instructions. Plates were read on the multiplexing imager Sector S600 (Meso Scale Discovery). All samples were measured in duplicate.

### Mucin glycosylation analysis

#### Isolation and purification of mucins secreted by Calu-3 cells

Cells were solubilized in 4 M guanidine chloride reduction buffer containing 10 mM DTT, 5 mM ethylenediaminetetraacetic acid, 10 mM benzamidine, 5 mM *N*-ethylmaleimide, 0.1 mg/ml soy bean trypsin inhibitor and 1 mM phenylmethanesulfonyl fluoride. Two ml of reduction buffer was added to each apical chamber and incubated overnight at room temperature. Cell suspensions were then gently agitated by pipetting and each of the 5 filter suspensions per condition were pooled in a single aliquot. CsCl was added to an initial density of 1.4 g/ml and mucins were purified by isopycnic density-gradient centrifugation (Beckman Coulter LE80 K ultracentrifuge; 70.1 Ti rotor, 417,600 g at 15°C for 72 hours). Fractions of 1 ml were collected from the bottom of the tube and analysed for periodic acid-Schiff (PAS) reactivity and density. The mucin-containing fractions were pooled, dialyzed against water and lyophilized.

#### Release of oligosaccharides from mucin by alkaline borohydride treatment

Mucins were submitted to β-elimination under reductive conditions (0.1 M KOH, 1 M KBH4 for 24 hours at 45°C) and the mixture of oligosaccharide alditols was dried on a rotavapor (Buchi) at 45°C. Borate salts were eliminated by several co-evaporations with methanol before purification by cation exchange chromatography (Dowex 50×2, 200-400 mesh, H + form).

#### Permethylation and mucin glycosylation analysis by MALDI TOF MS

Permethylation of the mixture of oligosaccharide alditols was carried out with the sodium hydroxide procedure described by Ciucanu and Kerek (1984) [45]. After derivatization, the reaction products were dissolved in 200 μl of methanol and further purified on a C18 Sep-Pak column (Waters, Milford, MA). Permethylated oligosaccharides were analyzed by MALDI TOF MS in a positive ion reflective mode as [M+Na]+. Quantification through the relative percentage of each oligosaccharide was calculated based on the integration of peaks on MS spectra.

### Statistics

Statistical analyses were performed using GraphPad Prism 8 Software. One-way analysis of variance (ANOVA) or Student *t* test were used in this study. P-values of p<0.05 were considered to indicate statistical significance. Each images presented in this work were representative images.

## ACKNOWLEDGEMENTS

We thank Nicolas Goudin of the Necker Institute imaging facility for his technical support. We thank Emmanuelle Bille for her careful reading of the manuscript. We thank Karine Bailly of the Cochin Institute Cytometry and Immunobiology facility (CYBIO) for her excellent technical assistance. This work was supported by ANR-15-CE15-0002-01 grants. XN and MC are also supported by INSERM, CNRS, Université de Paris and the Fondation pour la Recherche Médicale. CRM was supported by the Research Federation FRABio (Univ. Lille, CNRS, FR 3688, FRABio, Biochimie Structurale et Fonctionnelle des Assemblages Biomoléculaires), the CNRS (Unité mixte de Recherche CNRS/ULille 8576) and the Ministère de l’Enseignement Supérieur et de la Recherche.

---

**Supplementary figure 1: Calu-3 cells grown using AIC. (A)** Confocal 3D reconstructions of living Calu-3 cells showing accumulation of the mucus after two weeks of culture in AIC. The mucus was labeled with Alexa-conjugated Dextran (cyan) and cells were stained with Cell Trace Calcein Red Orange, AM (red). Bar: 20 μm. **(B)** Transmission electron microscopy images (transversal section). *: Tight junction; arrow: mucin-containing vesicles; μV: microvilli. M: mucus. Bar: 2 μm.

**Supplementary figure 2: Dying and living *N. meningitidis.*** Transmission electron microscopy images (longitudinal section) showing dying and living bacteria inside the mucus (A) and dying bacteria inside a cell (B). DB: Dead bacteria; LB: living bacteria. Bar: 1μm.

**Supplementary figure 3: Culture of Calu-3 cells in the presence of IL-4 and IL-13; and infection of Calu-3 cells. (A)** Measurement of the TEER of Calu-3 cell layer cultured in AIC using 0.4μm pore membrane or 3μm pore membrane, with or without addition of 5 ng/ml IL-4 or IL-13 for 24 hours. TEER were expressed as mean Ohm.cm2 ± SEM (statistical significance: *** p<0.001, ** p<0.01; One way-ANOVA; 0.4 Vs 3 μm pore membrane: three filters; Control Vs IL-4/13: 4 filters, two independent experiments). **(B)** Count of meningococci 24 hours after infection of Calu-3 cells, grown in AIC, with 10^2^ or 10^4^ or 10^6^ bacteria. Data were expressed as mean of CFU per filter ± SEM. Six filters, two independent experiments. **(C)** Confocal 3D reconstructions showing *N. meningitidis* proliferation. Twenty-four hours after infection of Calu-3 cells with 10^2^ or 10^4^ or 10^6^ bacteria, cells were fixed and immuno-stained with anti-2C4.3 antibody (green). Cells were stained using Alexa-conjugated phalloïdin (in red). Nuclei were stained with dapi (blue). Bar: 20 μm.

**Supplementary figure 4: Co-culture of *N. meningitidis* with *S. mitis* in broth**. BHI broth co-cultures were performed during 24 hours and *N. meningitidis* CFU determined. The number of meningococci after 24 hours of co-culture was expressed as mean percentage of the control experiment ± SEM (control: CFU of meningococci in mono-culture) (statistical significance: **** p<0.0001, * p<0.01; Student *t* test; Four wells, two independent experiments).

