## Supplementary figures and images for "Air-interfaced colonization model suggests a commensal-like interaction of *Neisseria meningitidis* with the epithelium, which benefit from colonization by *Streptococcus mitis*"

### Supplemental figures 1 to 4

SUPPLEMENTARY FIGURE 1

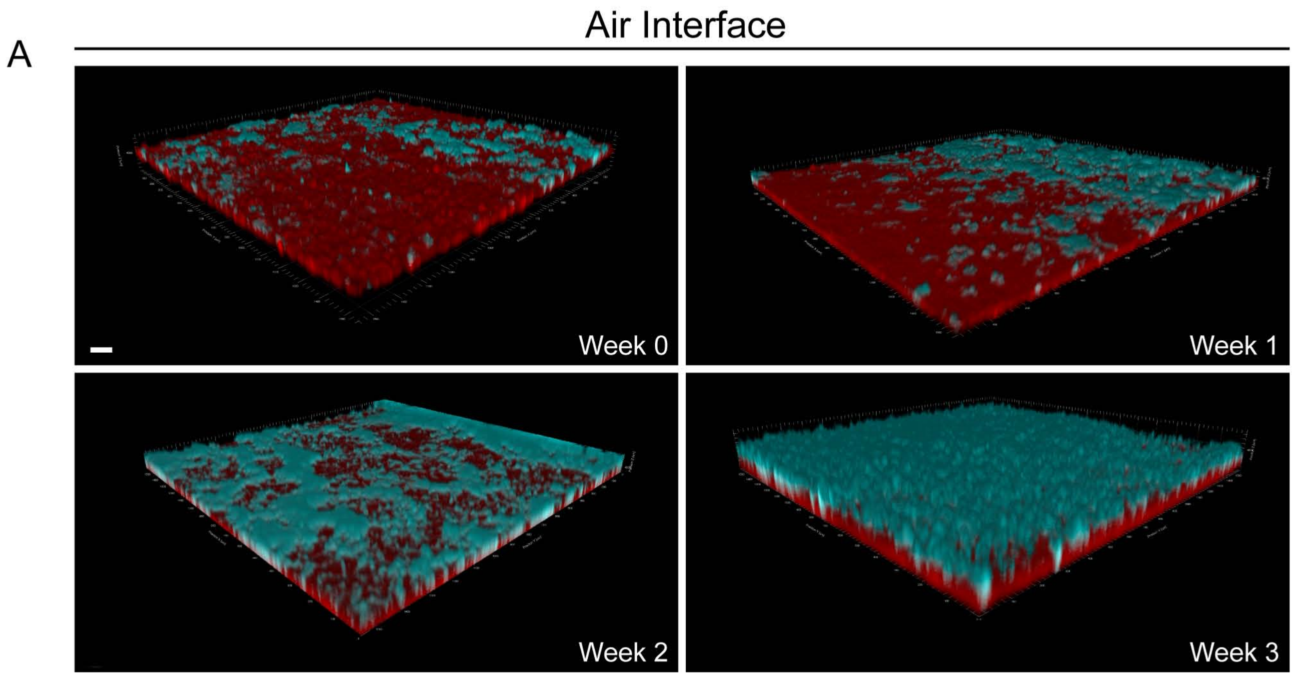


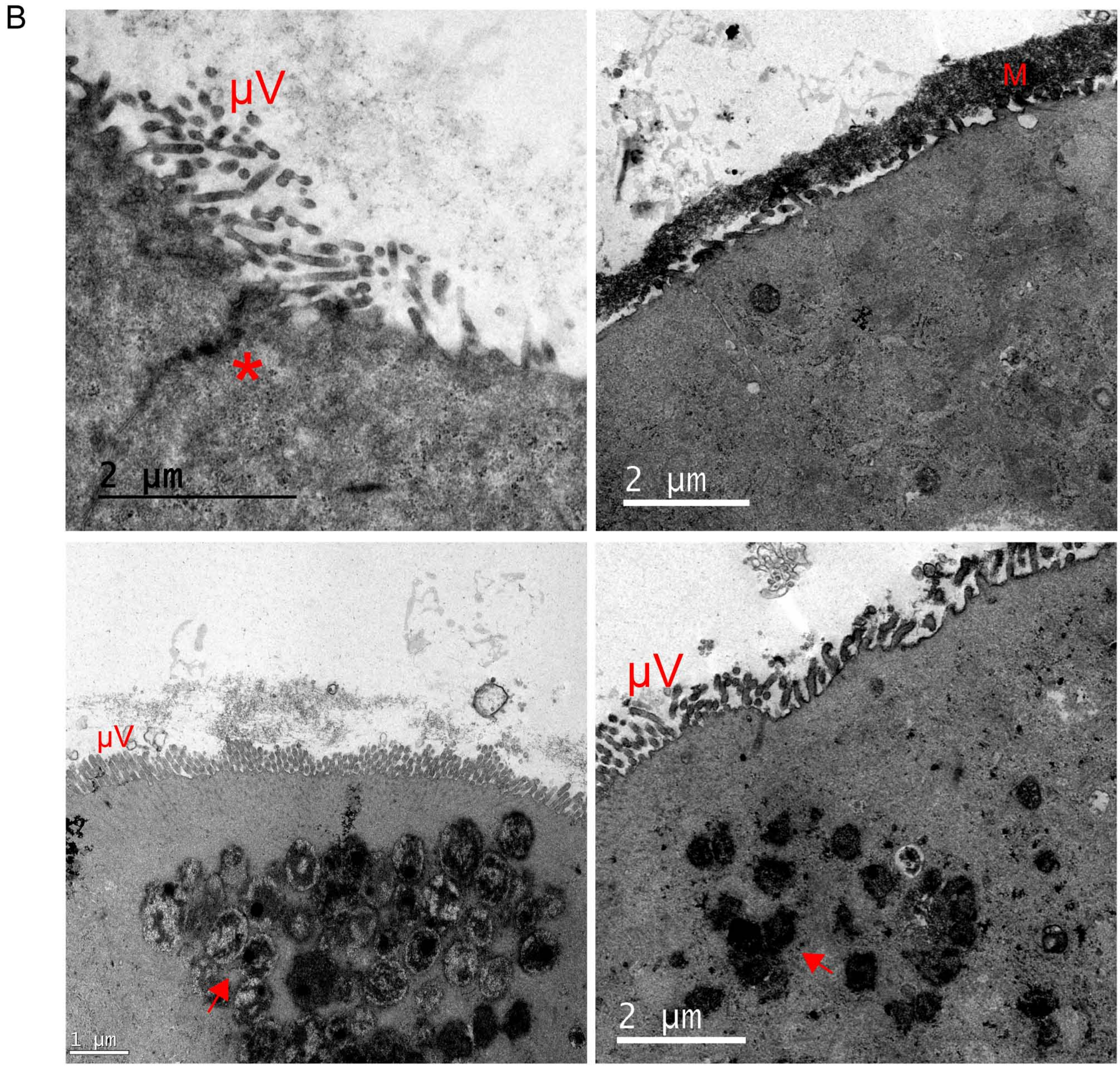

SUPPLEMENTARY FIGURE 2

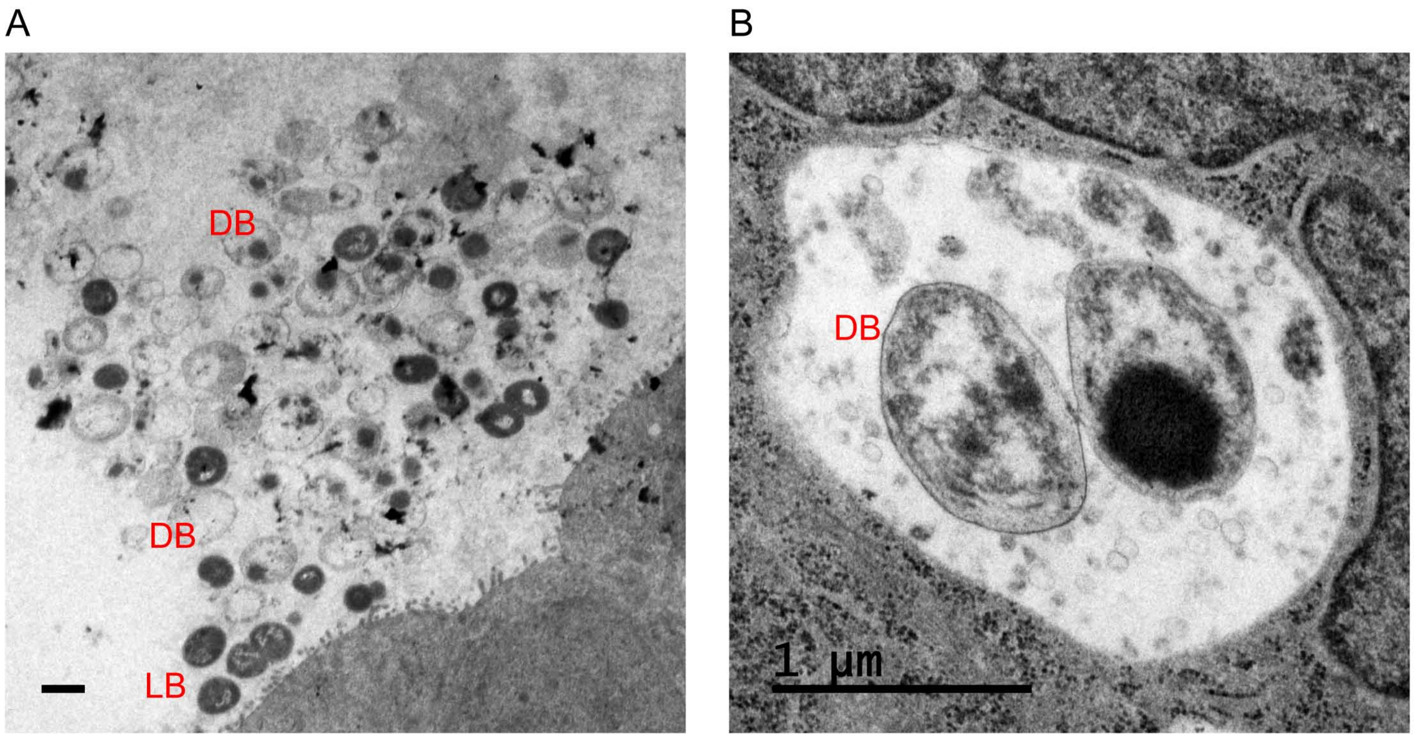

SUPPLEMENTARY FIGURE 3

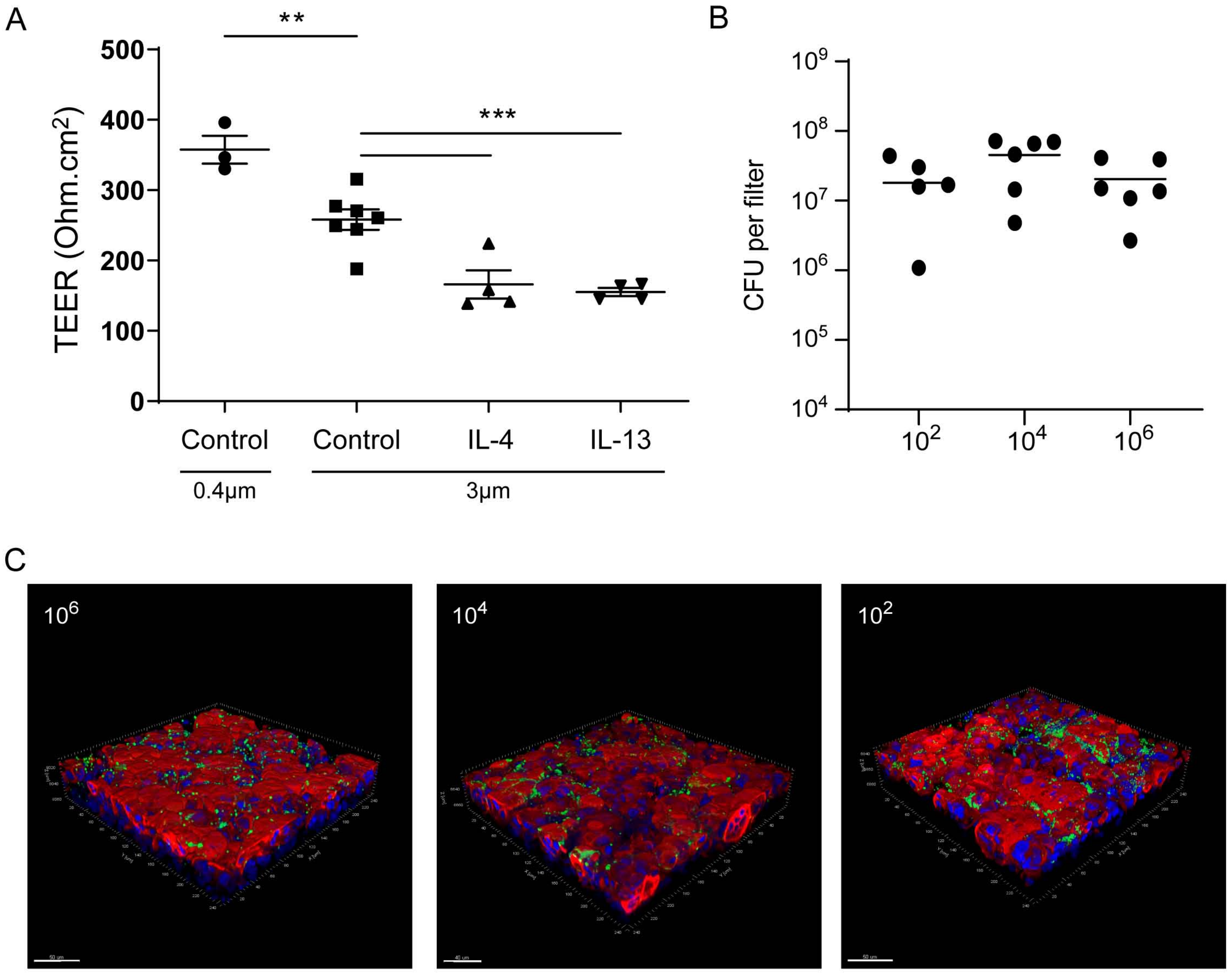

SUPPLEMENTARY FIGURE 4

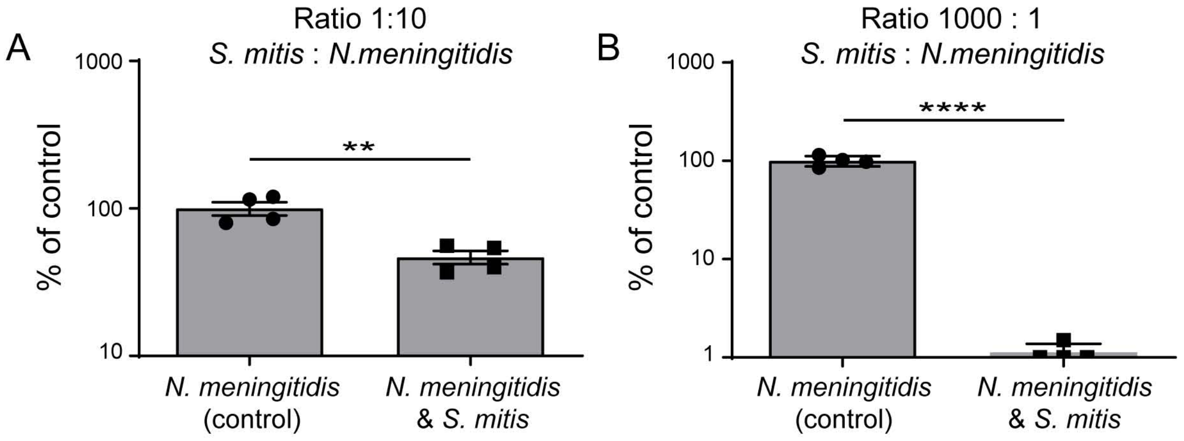
