## Supplemental Table S1 for "Air-interfaced colonization model suggests a commensal-like interaction of *Neisseria meningitidis* with the epithelium, which benefit from colonization by *Streptococcus mitis*"

**Table S1.** Oligosaccharides identified Calu-3 mucins. before (control) and after infection with *N. meningitidis* (*Nm*) or *S. mitis* (*Sm*) or heat-inactivated *S. mitis* (*Sm* HI) or co-infection with *N. meningitidis* and *S. mitis* (*Nm*+*Sm*)*.* The relative percentage of each oligosaccharide was calculated based on the integration of peaks on MS spectra.

^a^ mean ± SEM of each oligosaccharide. Two independent experiments of 5 different filters studied in bulk.

^b^ mean of each oligosaccharide. One experiment of 5 different filters studied in bulk.

| Proposed structures or sequences of oligosaccharides | [M+Na]^+^ | Calu-3 control^a^ | Calu-3 *Nm*^a^ | Calu-3 *Sm*^b^ | Calu-3 *Sm* HI^b^ | Calu-3 *Nm*+*Sm*^a^ |
| --- | --- | --- | --- | --- | --- | --- |
| 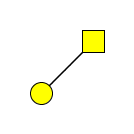 | 534 | 53.2± 1.2 | 34.4±1.6 | 82.2 | 56.3 | 86.7±8.8 |
| 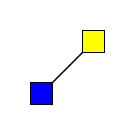 | 575 | 1.3± 1.2 | 0 | 0 | 0 | 0.3±0.3 |
| 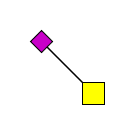 | 691 | 0.7±0.7 | 0 | 0 | 0 | 0 |
| 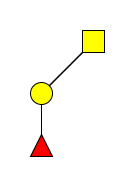 | 708 | 0.5±0.5 | 0 | 0 | 0 | 0 |
| 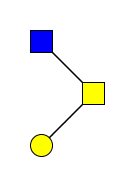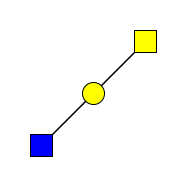 | 779 | 0.8 | 0.3±0 | 3.2 | 4 | 2.8±2.5 |
| 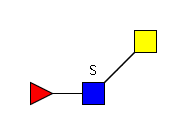 | 837 | 0 | 0.4±0.2 | 0 | 0 | 0 |
| 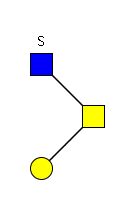 | 867 | 1.3±0.4 | 0.2±0.2 | 0 | 0 | 0 |
| 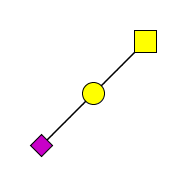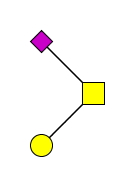 | 895 | 29.8±3.1 | 41.7±0.4 | 9.3 | 25.5 | 3±3 |
| 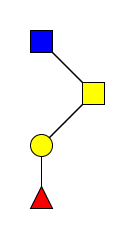 | 953 | 0.1±0.1 | 0 | 0 | 1.2 | 0 |
| 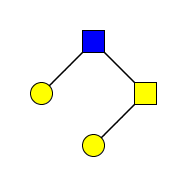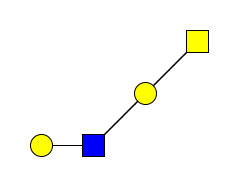 | 983 | 3.1±1.1 | 3.7±0.4 | 4.6 | 6.3 | 6.4±2.9 |
| 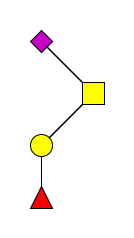 | 1069 | 0.2±0.2 | 0 | 0 | 0 | 0 |
| 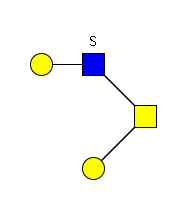 | 1071 | 0.3±0.1 | 0.2±0.2 | 0.4 | 1 | 0.8±0.8 |
| 1 Hex. 1 HexNAc. 1 NeuAc. GalNAcol | 1140 | 0.2±0.1 | 0.1±0.1 | 0 | 0.6 | 0 |
| 2 Hex. 1 HexNAc. 1 Fuc. GalNAcol | 1157 | 0.1±0.1 | 0 | 0 | 0 | 0 |
| 1 Hex. 2 HexNAc. 1 Fuc. GalNAcol | 1198 | 0 | 0 | 0 | 0 | 0 |
| 2 Hex. 2 HexNAc. GalNAcol | 1228 | 1.6±1.5 | 0.5±0.1 | 0 | 0.3 | 0 |
| 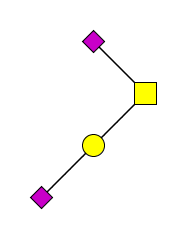 | 1256 | 3.6±1.6 | 13.4±1.2 | 0 | 3 | 0 |
| 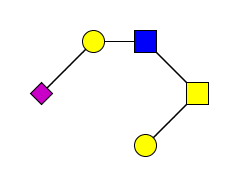 | 1344 | 1.2±0.5 | 4.2±0.4 | 0 | 1.3 | 0 |
| 3 Hex. 2 HexNAc. GalNAcol | 1432 | 0.4±0.3 | 0.4±0.1 | 0.2 | 0.3 | 0.2±0.1 |
| 3 Hex. 2 HexNAc. 1 SO3. GalNAcol | 1520 | 0.4±0.4 | 0.2±0.1 | 0 | 0.1 | 0 |
| 2 Hex. 1 HexNAc. 2 NeuAc. GalNAcol | 1705 | 1.3±1.3 | 0.6±0.1 | 0 | 0 | 0 |
| 3 Hex. 2 HexNAc. 1 NeuAc. GalNAcol | 1793 | 0.3±0.3 | 0.1±0.1 | 0 | 0 | 0 |
